# Generation and characterization of two immortalized dermal fibroblast cell lines from the spiny mouse (*Acomys*)

**DOI:** 10.1101/2022.12.23.521723

**Authors:** Michele N. Dill, Mohammad Tabatabaei, Manasi Kamat, Kari B. Basso, Chelsey S. Simmons

**Author notes:** Corresponding Author, (MND).

## Abstract

The spiny mouse (*Acomys*) is gaining popularity as a research organism due to its phenomenal regenerative capabilities. *Acomys* recovers from injuries to several organs without fibrosis. For example, *Acomys* heals full thickness skin injuries with rapid re-epithelialization of the wound and regeneration of hair follicles, sebaceous glands, erector pili muscles, adipocytes, and dermis without scarring. Understanding mechanisms of *Acomys* regeneration may uncover potential therapeutics for wound healing in humans. However, access to *Acomys* colonies is limited and primary fibroblasts can only be maintained in culture for a limited time. To address these obstacles, we generated immortalized *Acomys* dermal fibroblast cell lines using two methods: transfection with the SV40 large T antigen and spontaneous immortalization. The two cell lines (AcoSV40 and AcoSI-1) maintained the morphological and functional characteristics of primary *Acomys* fibroblasts, including maintenance of key fibroblast markers and ECM deposition. The availability of these cells will lower the barrier to working with *Acomys* as a model research organism, increasing the pace at which new discoveries to promote regeneration in humans can be made.

## Introduction

The spiny mouse (*Acomys*) is gaining popularity as a research organism, largely due to its phenomenal regenerative capabilities [1]. *Acomys* has been shown to regenerate damage to the skin [2–7], ear pinna [8–10], skeletal muscle [11], kidney [12], heart [13–15], and spinal cord [16,17]. Remarkably, all these findings have occurred within the last decade. The increased interest of *Acomys* as a research organism and its potential for future regenerative medicine applications creates a need for research tools that can be used to increase the accessibility of *Acomys* as a model organism and promote collaboration among researchers across different fields and universities.

However, the use of emerging research organisms has challenges. Many researchers do not have access to non-traditional animal facilities, and there are higher associated costs with maintaining non-traditional animal colonies. In the case of *Acomys*, the obstacles are even greater, as they are not sold by commonly used vendors like the Jackson laboratory and their colonies require maintenance procedures that differ from those of commonly housed rodents [1,18–20]. In addition, *Acomys* have longer gestation periods and smaller litters which makes building and maintaining a colony difficult. The limited presence and size of *Acomys* colonies available for research limits access to *Acomys* even as interest continues to increase.

Using primary cell cultures from *Acomys* carries similar challenges as primary cells require access to an *Acomys* colony, and repeated cell isolation can place stress on colonies being used for multiple projects. In addition, primary and secondary cell cultures can only be maintained *in vitro* for a short period of time before they enter replicative senescence and cell division is no longer possible [21,22]. Even prior to replicative senescence, primary and secondary cell cultures begin to show morphological and functional changes which limits their use to early passages. Isolation of primary cells is also time consuming, and the initial population of cells may be heterogeneous. While sorting for the cell type of interest is typically an option when working with primary isolates, the poor cross-reactivity of many antibodies presents even more challenges in *Acomys* [1,23,24].

In comparison to primary and secondary cell cultures, immortal cell lines have ‘infinite’ culturing capabilities and off-the-shelf availability, meaning they can easily be shared across research institutions. Immortal cell lines have acquired the ability to proliferate indefinitely through either artificial genetic modifications or spontaneous mutations. They are widely used because they are easy and inexpensive to maintain, manipulate, and expand. Immortal cells are a valuable tool for preliminary experiments because they are a pure population of cells which improves reproducibility. In addition, they obviate the need for consistent isolation of primary cells to support *in vitro* experiments.

Cell immortalization through artificial genetic modifications is often performed by transfection with viral oncogenes. Immortalization of cells through transfection with the simian virus 40 large T antigen (SV40-LT) has been used for decades and is described as a simple and reliable agent for the immortalization of many different cell types, and there are over 170 SV40-LT transfected cell lines available through ATCC alone [25–28]. Although the SV40-LT binds to many proteins, its interactions with the tumor suppressors retinoblastoma (Rb) and p53 are essential for bypassing replicative senescence [25,29]. Rb and p53 serve to prevent excessive cell growth and mutations by halting cell cycle progression, and their inhibition by SV40-LT promotes cell immortalization by preventing senescence. Cells transfected with SV40-LT also demonstrate stabilization of telomere length due to increased telomerase activity [30]. Viral transfection is not complete in all cells. Instead, within a cell culture, there are nonpermissive infections, semipermissive infections, and full transformations [31,32]. Fully transformed cells will eventually overtake the other populations in culture, but they can also be reliably isolated using selectable markers like antibiotic resistance genes, producing a culture of transformed cells.

Cells can also be immortalized through spontaneous mutations that result from prolonged subculture and genomic instability. Spontaneous immortalization of normal human cell lines is rare whereas normal rodent cells can regularly be established spontaneously [33,34]. One example is NIH3T3 fibroblasts which have been utilized in research since they were first developed by Todaro and Green in 1963 [35]. Loss of a key tumor suppressor gene like p53 is necessary for spontaneous immortalization, but another mutation such as chromosomal recombination or epigenetic silencing is also required [36,37]. Interestingly, the mutations required to escape replicative senescence can change with cell type and culture conditions [38–40], making it difficult to point to any specific mutations as being the key to spontaneous immortalization. Nonetheless, a variety of cell lines from different organisms including mice, humans, pigs, and chickens have been derived through spontaneous immortalization and utilized in experimentation [41–44].

To support the regenerative medicine community and to reduce the number of animals used in research, we sought to generate immortalized *Acomys* fibroblasts. We chose to focus on the dermis because regeneration of *Acomys* skin has been widely reported [2–7], but exact mechanisms remain elusive. Strikingly, *Acomys* can regenerate hairs, sebaceous glands, erector pili muscles, adipocytes, and the panniculus carnosus following full thickness excision wounds [2] and burn wounds [5] to the skin. We have chosen to focus on fibroblasts because we hypothesize that fibroblast activation plays a notable role in *Acomys* regeneration [45], and we are investing in *in vitro* experimental platforms to uncover specific regenerative mechanisms. We generated two *Acomys* dermal fibroblast lines—one through transfection with SV40-LT and one through extended subculture—and verified their functional similarity to primary *Acomys* fibroblasts (pAFs). These cell lines have been accepted into the ATCC general collection, and we expect availability in 2023.

## Materials and Methods

### Isolation of pAFs

Care and use of animals was conducted in accordance with the United States Department of Agriculture (USDA) and National Institutes of Health (NIH) guidelines and were approved by the UF Institutional Animal Care and Use Committee. *Acomys cahirinus* and CD-1 *Mus musculus* pups were obtained from UF breeding colonies.

*Acomys* pups were euthanized within 3 days of birth and the dorsal skin was removed. The tissue was incubated overnight in 0.2% dispase II (Roche) in DMEM at 4°C, after which the dermis and epidermis were separated with forceps. The dermis was dissociated by incubation in 0.24% collagenase type I (Gibco) in PBS at 37°C for 60-90 minutes with constant agitation. A cell suspension was obtained using a vacuum filtered conical tube, and the cells were rinsed and cultured in DMEM/F:12 supplemented with 10% fetal bovine serum (Gibco), 10% NuSerum (Corning), 1% Gentamicin/Amphotericin B (Gibco), and 0.1% insulin-transferrin-selenium (Gibco). Isolation of cells from CD-1 *Mus musculus* pups followed the same procedure.

### SV40LT transfection of pAFs

Fibroblasts were isolated from an *Acomys* neonate and expanded for two passages on soft silicone (Sy1gard^™^ 527 Silicone Dielectric Gel, Dow Inc.) coated with 0.01mg/mL rat tail collagen I (Corning) prior to being frozen and shipped to ALSTEM, Inc for immortalization. At ALSTEM, cells were cultured in provided collagen-coated Sy1gard 527 well plates and using the media formulation described above. Sy1gard 527 was used to mimic the stiffness of *in vivo* tissue and avoid phenotypic changes in the pAFs during culture. 100,000 cells were infected with a lentivirus encoding the SV40 large T antigen and puromycin N-acetyltransferase. The cells were selected by puromycin at 2ug/mL and passaged for 2-3 passages. The expression of SV40 and puromycin resistance genes was confirmed via PCR (S1 Fig) before the cells were frozen.

Upon receiving the cells, continuous proliferation was confirmed by passaging the cells at a constant interval (3 days) and seeding density (5,300 cells/cm^2^). Growth curves were generated by calculating the population doubling (PDL) at each passage using the formula PDL = PDL_0_ + 3.32(LogC_f_ — LogC_0_) where C_f_ is the final cell count at the time of passaging and C_0_ is the seeding number. The cells were intermittently frozen so proliferation could be assessed across both passages and freeze/thaw cycles. We utilized the media formulation that has been optimized for our pAFs (described above) for both immortalized lines to directly compare them to pAFs. However, we have demonstrated that both lines also retain continuous proliferation in the presence of a more commonly used formulation (S2 Fig).

### Spontaneous immortalization of primary Acomys fibroblasts

Fibroblasts were isolated from three *Acomys* neonates from separate litters and cultured until logarithmic growth was re-established following a period of reduced proliferation. Cells were passaged at 80% confluency, and a constant seeding density (5,300 cells/cm^2^) was used. The cells were classified as immortalized once proliferation increased for 3 passages following the period of reduced proliferation, or crisis.

One set of immortalized *Acomys* fibroblasts (AcoSI-1) was chosen for further characterization. Continuous proliferation of AcoSI-1 cells was confirmed by passaging the cells at a constant interval (3 days) and seeding density (5,300 cells/cm^2^). Growth curves were generated by calculating the PDL at each passage as described above. The cells were intermittently frozen so proliferation could be assessed across both passages and freeze/thaw cycles.

### Assessment of fibroblast markers via western blot

Cell lysates were prepared in RIPA buffer, and equal amounts of protein were loaded into NuPage 4-12% Bis-Tris Midi gels (Invitrogen) submerged in NuPage MES SDS Running Buffer (Life Technologies). The proteins were transferred onto a nitrocellulose membrane using the SureLock^™^ Tandem Midi Blot Module (Life Technologies) and NuPage Transfer Buffer (Life Technologies) containing 10% methanol. The membrane was blocked with 5% powdered milk in tris-buffered saline for 1 hour at room temperature. Membranes were incubated with either a mouse monoclonal anti-alpha smooth muscle actin (Abcam ab7817, 0.341ug/mL) or a rabbit monoclonal anti-vimentin antibody (Abcam ab92547, 1:5000 dilution) with a mouse monoclonal anti-GAPDH loading control (Arigo biolaboratories ARG10112, 1:5000 dilution) overnight at 4°C. Incubation with HRP secondary antibodies (Arigo biolaboratories ARG65350, 1:5000 dilution or enQuire BioReagents QAB10303, 1:15,000 dilution) was performed for 1 hour at room temperature. Signal was produced using SuperSignal^™^ West Pico PLUS Chemiluminescent Substrate (Thermo Scientific) and the blots were imaged on a LI-COR Fc Imager (Odyssey).

### Assessment of contractility with traction force microscopy

Cell contractility was evaluated by using traction force microscopy (TFM) as described previously [46]. Cells were seeded on 8kPa polyacrylamide hydrogels coated with fluorescent nano-beads (Cell&Soft) and allowed to adhere for 12 hours. The samples were then mounted on a Nikon microscope with a 37°C chamber, and DMEM media was replaced with CO_2_-independent media (Leibovitz) prior to the test. Fluorescent images of the nano-beads were captured before and after cell detachment with a 1% Triton-X and 200 mM KOH solution. Bead displacement was quantified and used to calculate the root-mean-squared values of stress and strain energy. For each group of tests, at least 50 cells were measured and results were reported as the mean ± standard deviation. The results were compared using one-way analysis of variance (ANOVA) with the Bonferroni post hoc test. p-value < 0.05 was considered statistically significant.

### Preparation of cell derived matrices

Cell derived matrices (CDMs) were obtained by seeding pAFs, AcoSV40, or AcoSI-1 fibroblasts onto collagen-functionalized Sy1gard 527 PDMS. Primary *Mus* fibroblasts and NIH3T3 fibroblasts were included as controls, and 3 sets of CDMs were made from each cell type (15 samples total). Prior to cell seeding, the substrate was plasma treated to oxidize the surface before functionalizing with (3-Aminopropyl)trimethoxysilane, followed by glutaraldehyde and then rat tail collagen I. Functionalization with collagen I is needed to prolong adherence of *Acomys* cells to the substrate. Cells were cultured for 7-28 days in media containing 25mg/mL Ficoll 400, with immortalized cells requiring less time in culture to generate similar protein mass as primary cells. Following this period, CDMs were decellularized by treating with a solution of 0.5%Triton X-100 and 0.3M ammonium hydroxide in PBS for 5 minutes. The presence of residual DNA was reduced by treating decellularized CDMs with 10ug/mL DNase I at 37oC for 30 minutes.

Decellularized CDMs were homogenized with RIPA Buffer and a tissue homogenizer (Fisherbrand^™^ 150 Handheld Homogenizer). The samples were centrifuged at 3200g and 4°C for 15 minutes, followed by 14,000g for 2 minutes. The supernatant was removed and stored at - 20°C as the RIPA soluble protein fraction. The pellet was resuspended in a membrane solubilization buffer containing 40mM Tris-Cl (pH 8.0), 7M urea, 2M thiourea, 0.25% w/v ASB-14, and 0.25% NP-40 and incubated at room temperature for 15 minutes. The samples were vortexed for 1 minute, then centrifuged at 3200g for 30 minutes. The supernatant was removed and diluted by 2.5 with dH_2_O and stored at −20°C as the RIPA insoluble protein fraction. Prior to analysis, both the RIPA soluble and RIPA insoluble protein fractions were precipitated in ice-cold acetone overnight. The solutions were centrifuged at 3200g and 4°C for 15 minutes. The acetone was fully removed, and the remaining pellets were resuspended in 2M urea. Equal concentrations of the RIPA soluble and RIPA insoluble protein fractions from each sample were combined, and label-free quantitative proteomics was performed by the UF Mass Spectrometry Research and Education Center.

### Label-free quantitative proteomics of CDM samples

#### In Solution Digestion

Total protein was determined on a Qubit and the appropriate volume of each sample was taken to equal 20 μg total protein for digestion. The samples were digested with sequencing grade trypsin/lys C enzyme (Promega) using manufacture recommended protocol. The samples were diluted with 50mM ammonium bicarbonate buffer. The samples were incubated at 56°C with 1.0 μL of dithiothreitol (DTT) solution (0.1 M in 50 mM ammonium bicarbonate) for 30 minutes prior to the addition of 3.0 μL of 55 mM iodoacetamide in 50 mM ammonium bicarbonate. Samples with iodoacetamide were incubated at room temperature in the dark for 30 min. The trypsin/lys C was prepared fresh as 1 μg/μL in reconstitution buffer (provided with the enzyme). 1 μL of enzyme was added and the samples were incubated at 37°C overnight. The digestion was stopped with addition of 0.5% trifluoracetic acid. The MS analysis is immediately performed to ensure high quality tryptic peptides with minimal non-specific cleavage.

#### Q Exactive HF Orbitrap

Nano-liquid chromatography tandem mass spectrometry (Nano-LC/MS/MS) was performed on a Thermo Scientific Q Exactive HF Orbitrap mass spectrometer equipped with an EASY Spray nanospray source (Thermo Scientific) operated in positive ion mode. The LC system was an UltiMate^™^ 3000 RSLCnano system from Thermo Scientific. The mobile phase A was water containing 0.1% formic acid and the mobile phase B was acetonitrile with 0.1% formic acid. The mobile phase A for the loading pump was water containing 0.1% trifluoracetic acid. 5 mL of sample is injected on to a PharmaFluidics mPAC^TM^ C18 trapping column (C18, 5 μm pillar diameter, 10 mm length, 2.5 μm inter-pillar distance). at 25 μl/min flow rate. This was held for 3 minutes and washed with 1% B to desalt and concentrate the peptides. The injector port was switched to inject, and the peptides were eluted off of the trap onto the column. PharmaFluidics 50 cm mPAC^TM^ was used for chromatographic separations (C18, 5 μm pillar diameter, 50 cm length, 2.5 μm inter-pillar distance). The column temperature was maintained 40°C. A flow-rate of 750 nL/min was used for the first 15 minutes and then the flow was reduced to 300 nL/min. Peptides were eluted directly off the column into the Q Exactive system using a gradient of 1% B to 20%B over 100 minutes and then to 45%B in 20 minutes for a total run time of 150 minutes.

The MS/MS was acquired according to standard conditions established in the lab. The EASY Spray source operated with a spray voltage of 1.5 KV and a capillary temperature of 200°C. The scan sequence of the mass spectrometer was based on the original TopTen^™^ method; the analysis was programmed for a full scan recorded between 375 – 1575 Da at 60,000 resolution, and a MS/MS scan at resolution 15,000 to generate product ion spectra to determine amino acid sequence in consecutive instrument scans of the fifteen most abundant peaks in the spectrum. The AGC Target ion number was set at 3e6 ions for full scan and 2e5 ions for MS^2^ mode. Maximum ion injection time was set at 50 ms for full scan and 55 ms for MS^2^ mode. Micro scan number was set at 1 for both full scan and MS^2^ scan. The HCD fragmentation energy (N)CE/stepped NCE was set to 28 and an isolation window of 4 *m/z*. Singly charged ions were excluded from MS^2^. Dynamic exclusion was enabled with a repeat count of 1 within 15 seconds and to exclude isotopes. A Siloxane background peak at 445.12003 was used as the internal lock mass.

HeLa protein digest standard is used to evaluate the integrity and the performance of the columns and mass spectrometer. If the number of protein ID’s from the HeLa standard falls below 2700, the instrument is cleaned and new columns are installed.

All MS/MS spectra were analyzed using Sequest (Thermo Fisher Scientific; version IseNode in Proteome Discoverer 2.4.0.305). Sequest was set up to search *Mus saxicola* Tax ID: 10094 assuming the digestion enzyme trypsin. Sequest was searched with a fragment ion mass tolerance of 0.020 Da and a parent ion tolerance of 10.0 ppm. Carbamidomethyl of cysteine was specified in Sequest as a fixed modification. Met-loss of methionine, met-loss+Acetyl of methionine, oxidation of methionine and acetyl of the n-terminus were specified in Sequest as variable modifications.

Precursor ion intensity label free quantitation was done using Proteome Discoverer (Thermo Fisher Scientific vs 2.4.0.305). The two groups (B33p4 vs Hp4) were compared using a “non-nested” study factor. Normalization was derived by using all peptides. Protein abundances were calculated by summed abundances, meaning the protein abundances are calculated by summing sample abundances of the connected peptide groups. Fisher’s exact test (pairwise ratio-based) was used to calculate p-values with no missing value imputation included. Adjusted p-values were calculated using Benjamini-Hochberg.

### Determining sex of immortalized Acomys fibroblast lines

Genomic DNA was isolated from 1 million cells of each cell line using the QIAamp®DNA Micro Kit (QIAGEN) following the manufacturer’s guidelines. Control gDNA was obtained from the peripheral blood of an adult male *Acomys*. DNA quantity was determined using a BioTek Synergy H1 plate reader (Agilent Technologies), and PCR reactions were performed using *Acomys-specific* primers for the target SrY. The PCR products were added to 2% E-Gel agarose gels with SYBR Safe DNA Stain (Invitrogen) and run in an E-Gel PowerSnap system (Invitrogen). Images were acquired using a LI-COR Fc Imager (Odyssey).

## Results

### Immortalized *Acomys* fibroblasts demonstrate continuous proliferation

Primary *Acomys* fibroblasts (pAFs) were isolated, cultured on silicone-coated tissue culture plates (Sy1gard 527, Elastic Modulus ~5 kPa), and sent to ALSTEM, Inc for immortalization with the SV40 large T antigen via lentiviral infection. The cells were selected by puromycin, and successful immortalization was confirmed through PCR assessment of SV40 and puromycin resistance genes (S1 Fig). We refer to these cells as “AcoSV40” cells. Continuous proliferation was confirmed by passaging the cells at a constant interval (3 days) and seeding density (5,300 cells/cm^2^) and calculating the population-doubling level at each passage. The AcoSV40 cells demonstrated continuous proliferation over 30 passages and multiple freeze/thaw cycles (Fig 1a).

**Figure 1.**
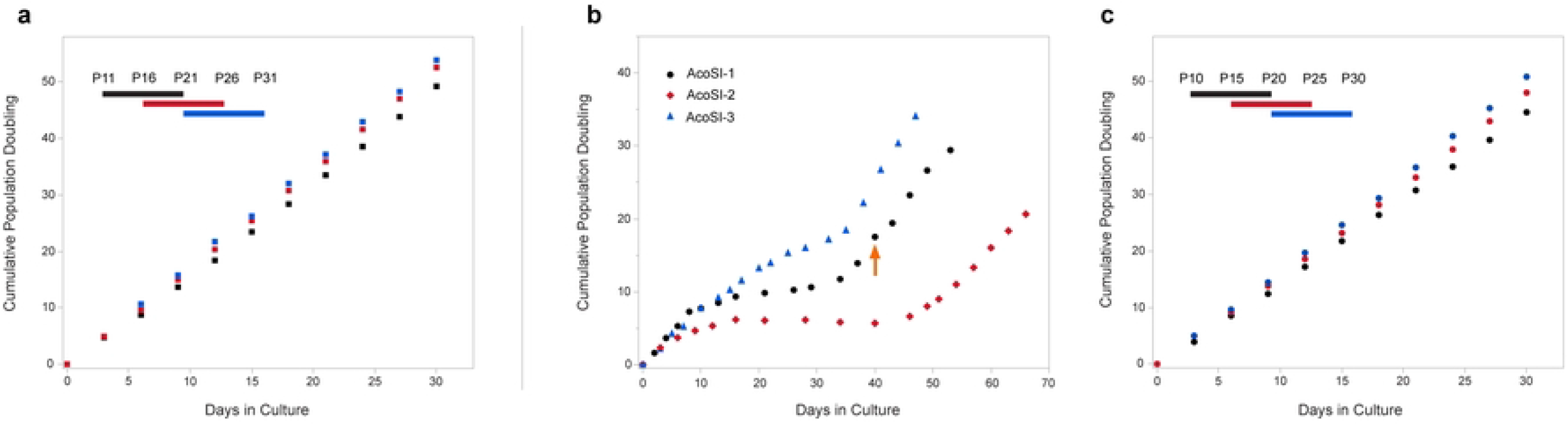
Immortalized AcoSV40 and AcoSI-1 fibroblasts demonstrate logarithmic growth over at least 30 passages. (a) AcoSV40 fibroblasts maintain logarithmic growth across 30 passages and 3 freeze/thaw cycles. (b) pAFs subjected to continuous subculture spontaneously immortalize after about 40 days. (c) AcoSI-1 fibroblasts maintain logarithmic growth across 30 passages and 3 freeze/thaw cycles.

Spontaneous immortalization of pAFs was performed through extended subculturing where the seeding density was kept constant (5,300 cells/cm^2^), and the cells were passaged upon reaching 80-90% confluency. As shown in Fig 1b, the fibroblasts demonstrated consistent proliferation for several passages prior to experiencing a period of ‘crisis’ during which there was a progressive decline in proliferation. A subpopulation of these cells eventually recovered and began to proliferate, overtaking the culture. Once this occurred, the cells were deemed immortalized and were named “AcoSI” cells. pAFs were isolated from three different animals, and the process was repeated to demonstrate replicability. AcoSI cells were generated within 13-17 generations and 40-55 days in culture, similar to reports by Todero and Green, in which mouse endothelial fibroblasts exited crisis after 15-30 generations and 45-75 days in culture [35]. The decreased time to immortalization in *Acomys* fibroblasts may be due in part to higher rates of proliferation in *Acomys* fibroblast cultures when compared to mouse fibroblast cultures (S3 Fig).

AcoSI-1 fibroblasts were chosen for futher assays due to the marked crisis period and median proliferation rates compared to the other two AcoSI lines. The passage at which the *Acomys* fibroblasts were deemed to be immortalized is marked by the arrow in Fig 1b and was recorded as AcoSI-1 passage 1. The same procedure was followed to determine continuous proliferation in the AcoSI-1 cells as the AcoSV40 cells. The cells were passaged every three days and seeded at a constant cell density. Population doubling remained consistent over 30 passages and multiple freeze/thaw cycles (Fig 1c).

### AcoSV40 and AcoSI fibroblasts retain similar morphological characteristics compared to pAFs

The morphology of the two immortalized *Acomys* fibroblast lines was compared to passage 1 pAFs via phase imaging. As depicted in Fig 2a, all three cell types share a similar size and shape, with the immortal cell lines appearing less spread than the pAFs and more cuboidal in shape.

**Figure 2.**
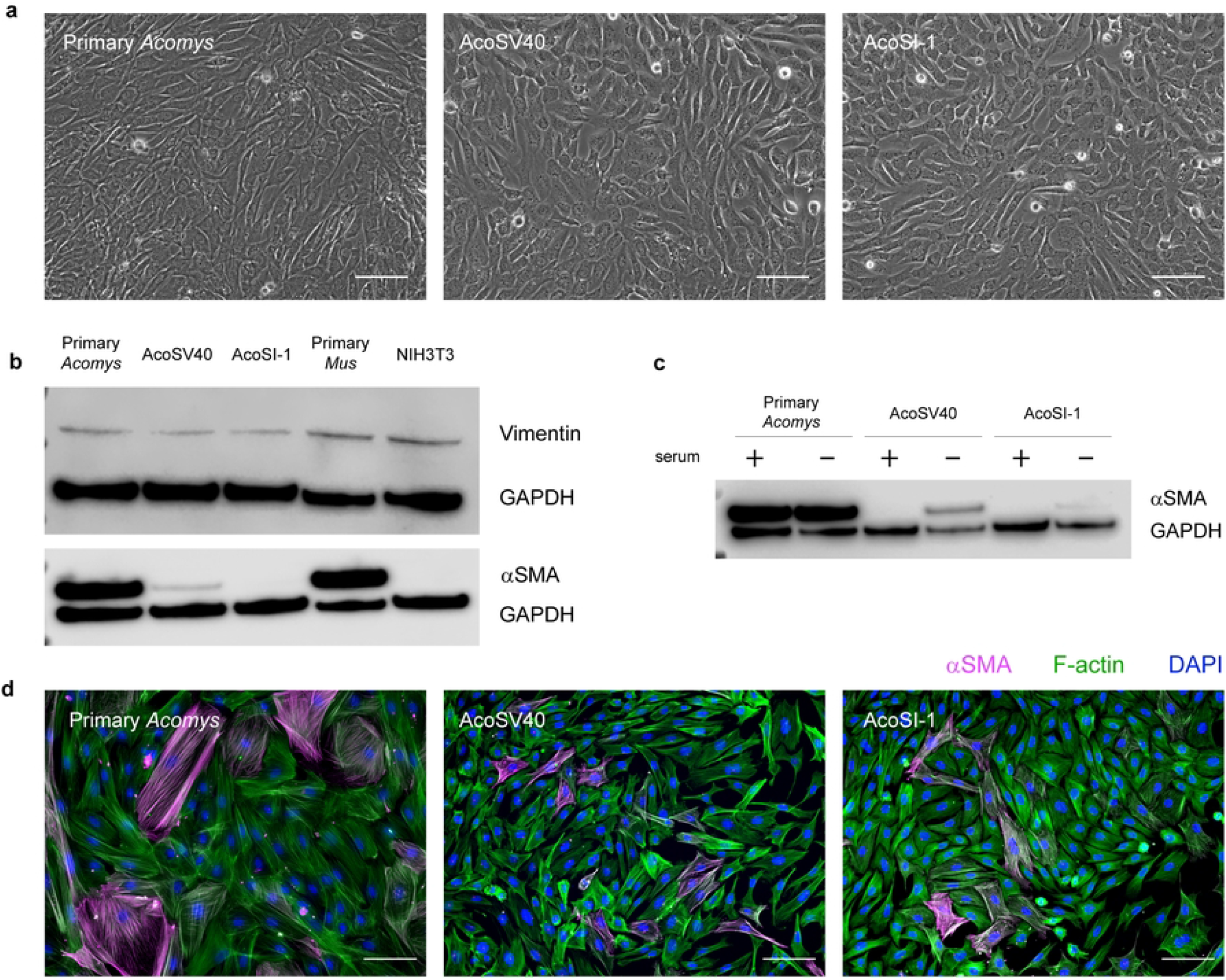
AcoSV40 and AcoSI-1 fibroblasts maintain morphological characteristics and fibroblast markers (vimentin and αSMA) of pAFs. (a) Phase images of passage 1 pAFs, AcoSV40, and AcoSI-1 fibroblast demonstrate similar morphology; scale = 100um. (b) pAFs, AcoSV40, and AcoSI-1 fibroblasts express vimentin at similar levels, but αSMA is weekly present in AcoSV40 and absent in AcoSI-1 fibroblasts under standard culture conditions. We demonstrated that under serum starvation, AcoSV40 and AcoSI-1 can (c) upregulate αSMA and (d) co-localize it to stress fibers; scale = 100μm.

The AcoSV40 and AcoSI lines were also assessed for the fibroblast markers vimentin and alpha smooth muscle actin (αSMA) to confirm relevant markers are retained during immortalization. Primary *Mus musculus* and immortalized *Mus musculus* (NIH3T3) fibroblasts were included as a comparison. As demonstrated by western blot (Fig 2b), all cell types maintain the vimentin marker and express it at similar levels under normal culture conditions. The AcoSV40 fibroblasts express αSMA at low levels in normal culture conditions compared to pAFs. Interestingly, the AcoSI-1 cells did not produce αSMA bands when assessed via western bot. However, this finding was shared with the NIH3T3 cells, which are also a spontaneously immortalized cell line. NIH3T3 fibroblasts are known to upregulate αSMA in culture when treated with TGF-β1 [47–49], suggesting that AcoSI-1 fibroblasts may also be able to upregulate αSMA under different culture conditions. TGF-β1 was not a good candidate for our purposes due to the lack of response documented in *Acomys* fibroblasts [50]. Instead, we assessed αSMA upregulation under serum starvation conditions and found that both AcoSV40 and AcoSI-1 fibroblasts produce more αSMA in the absence of serum, although AcoSI-1 cells make only a small quantity (Fig 2c). Both AcoSV40 and AcoSI-1 cells were also able to co-localize αSMA to stress fibers (Fig 2d), supporting their utility in fibroblast activation studies.

### AcoSV40 and AcoSI-1 fibroblasts retain functional characteristics of pAFs

Contractility and deposition of extracellular matrix (ECM) are important functional characteristics of fibroblasts and play a key role in stabilization of the wound bed following injury. We assessed the contractility of AcoSV40 and AcoSI-1 fibroblasts through traction force microscopy and found that they generate similar average (root-mean-square traction, rmst) and maximum (Max rmst) traction forces as pAFs (Fig 3). Strain energy (integrated traction force and deformation over the area of the cell) is unsurprisingly lower for immortalized fibroblasts since they have a smaller spread area than primary cells. Traction force microscopy confirms maintenance of contractile function in both AcoSV40 and AcoSI-1 immortalized lines.

**Figure 3.**
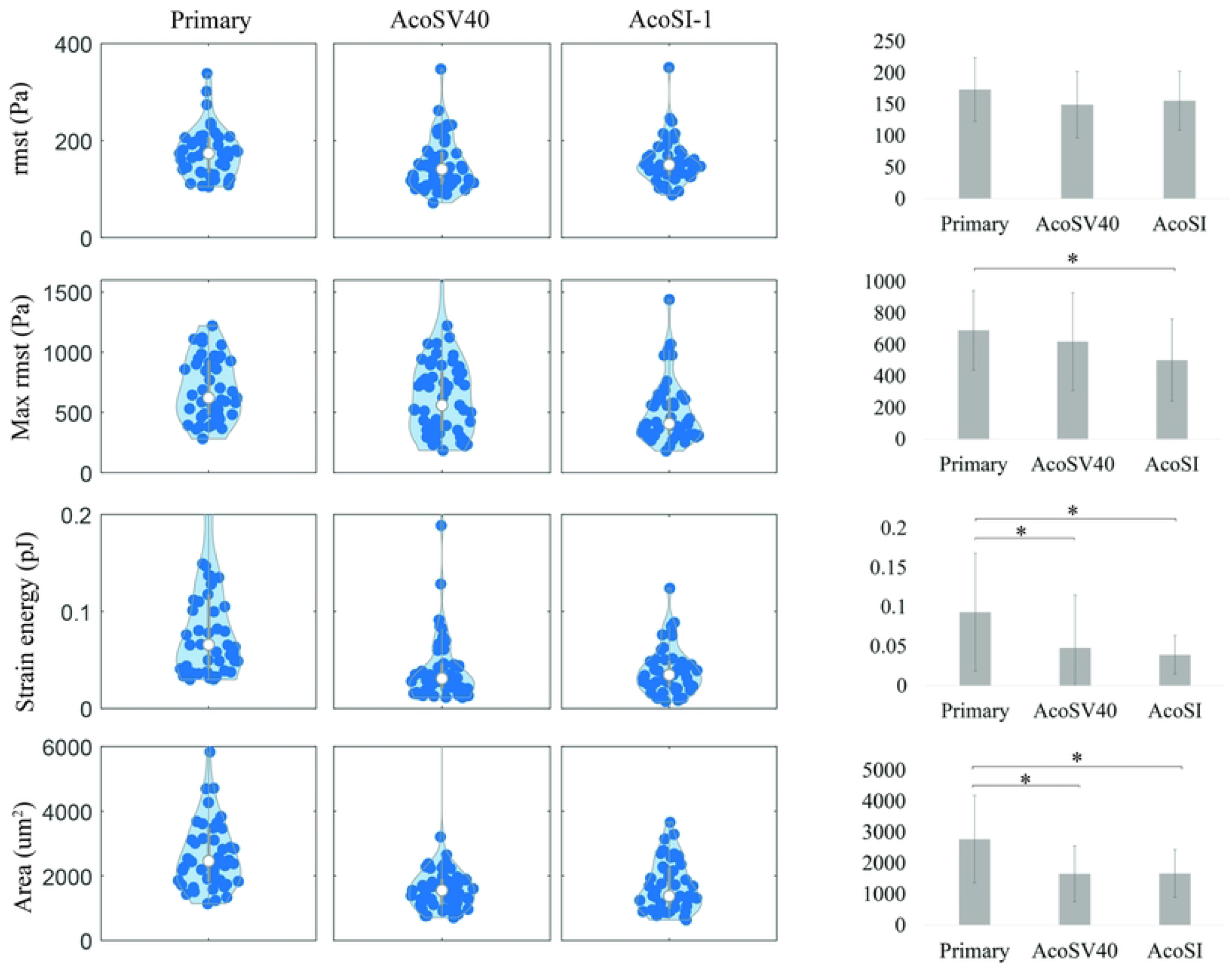
Traction forces of pAFs, AcoSV40, and AcoSI-1 fibroblasts are similar, and differences in strain energy correlate with differences in cell area. Cells were seeded on 8kPa polyacrylamide gels to evaluate root-mean-square of traction (rmst), max rmst, strain energy, and area. pAFs have higher maximum contraction stresses than AcoSI-1 cells, but similar average stresses to both immortalized lines. While pAFs have higher strain energy (i.e., do more work) than the immortalized fibroblasts, it is likely a function of their larger area. *p<0.05 as assessed by ANOVA with Bonferroni test.

Fibroblasts are responsible for maintaining the ECM proteome, composed of hundreds of proteins and proteoglycans which are encoded by over 1000 genes [51]. To capture this complexity, we compared ECM production in primary and immortalized *Acomys* fibroblasts by generating cell derived matrices (CDMs). CDMs recapitulate native tissue structure and composition *in vitro* and are useful for studying bulk matrix composition in a controlled setting. CDMs were generated in culture for 1-4 weeks depending on cell type, homogenized, and analyzed via label-free quantitative proteomics. No significant differences in CDM composition were found between pAFs and AcoSV40 fibroblasts, while AcoSI-1 CDMs shared 88% of proteins identified with pAFs (Fig 4a). The proteins that differed between the two samples were involved in biological processes related to metabolism and translation, based on a GO enrichment analysis. In comparison, NIH3T3s, a commonly used substitute for Mus musculus cells, only shared 75% of proteins with their counterpart (Fig 4b). Based on these results, AcoSV40 and AcoSI-1 are representative of pAFs in experiments related to ECM deposition.

**Figure 4.**
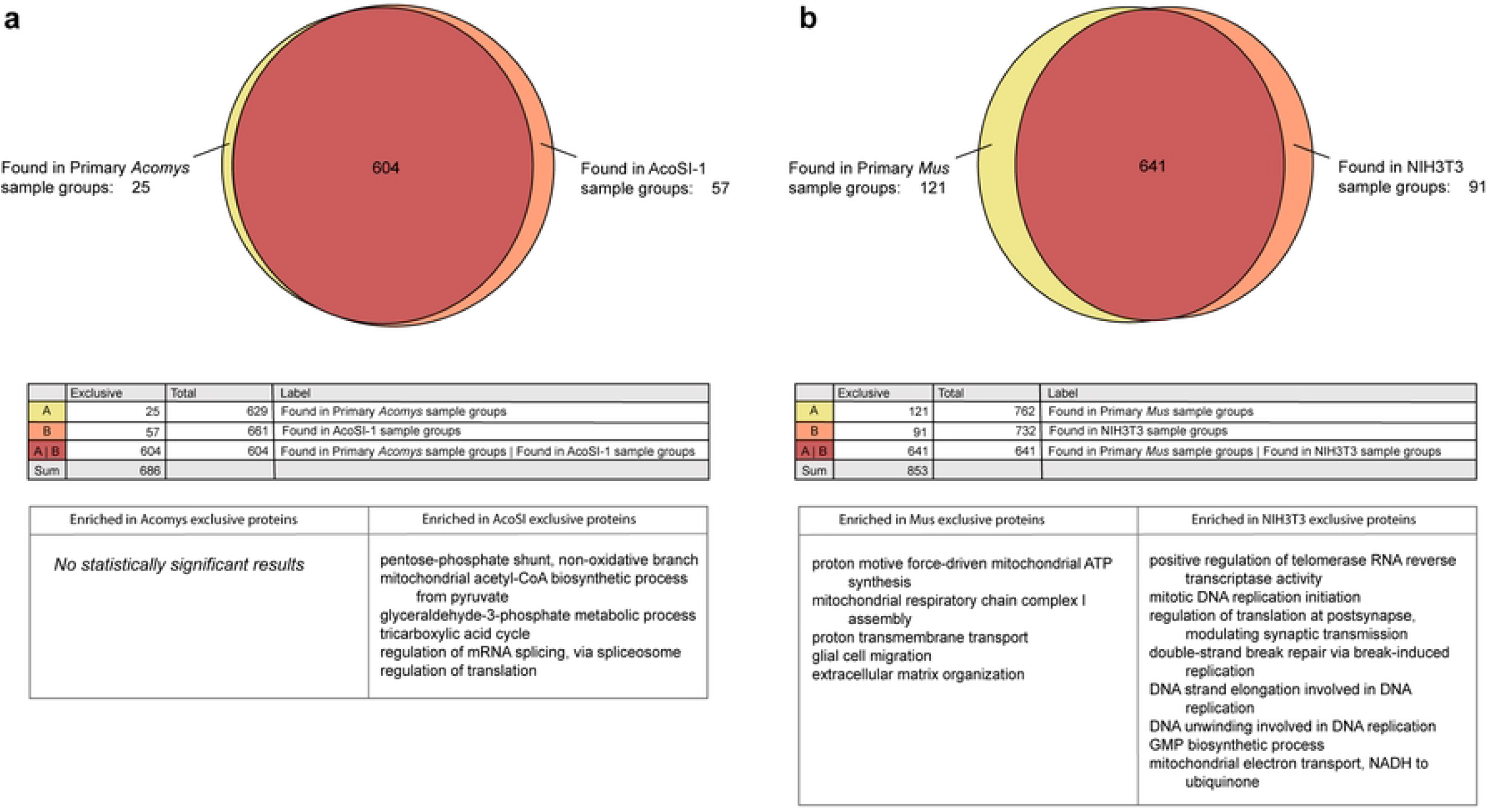
Immortalized lines AcoSV40 and AcoSI-1 share most proteins with pAFs. The comparison of AcoSV40 and pAF CDMs via mass spectrometry resulted in no significant difference in deposited proteins. (a) AcoSI-1 CDMs shared 604 out of 686 proteins identified with pAFs and enriched biological processes within the dissimilar proteins were unrelated to ECM organization. (b) In comparison, NIH3T3 fibroblasts share 641 out of 853 identified proteins with Mus primary fibroblasts, some of which are related to ECM organization. GO biological processes were determined by running the proteins that were exclusive to Primary *Acomys*, AcoSI-1, Primary *Mus*, and NIH3T3 fibroblasts through the PANTHER classification system. For brevity, only the top eight enriched biological processes were reported in the NIH3T3 exclusive proteins, all of which had a fold enrichment change of over 50.

### Determining sex of immortalized Acomys cell lines

It is becoming increasingly evident that considering the sex of cell lines is important. Genes expressed on the sex chromosomes can have an impact on a cell’s biology, and cells differ according to sex, regardless of their exposure to sex hormones. The basis of differences between male and female cells and examples of known differences between male and female cells from a variety of tissues are reviewed by Shah et al [52]. Since we were unable to sex the *Acomys* pups from which the cells were isolated, we determined the sex of our immortalized lines via PCR using primers designed for the *Acomys* SRY gene. The SRY gene provides instructions for making the sex-determining region Y protein which is located on the Y chromosome and is involved in male-typical sex development. Genomic DNA was obtained from cell cultures of the four immortalized lines and the blood of a male *Acomys* and assessed for the presence of SRY (Fig 5a). DNA from AcoSV40 fibroblasts did not contain the SrY gene, while all AcoSI cell lies did, meaning AcoSV40 cells are female while all AcoSI cells are male (Fig 5b).

**Figure 5.**
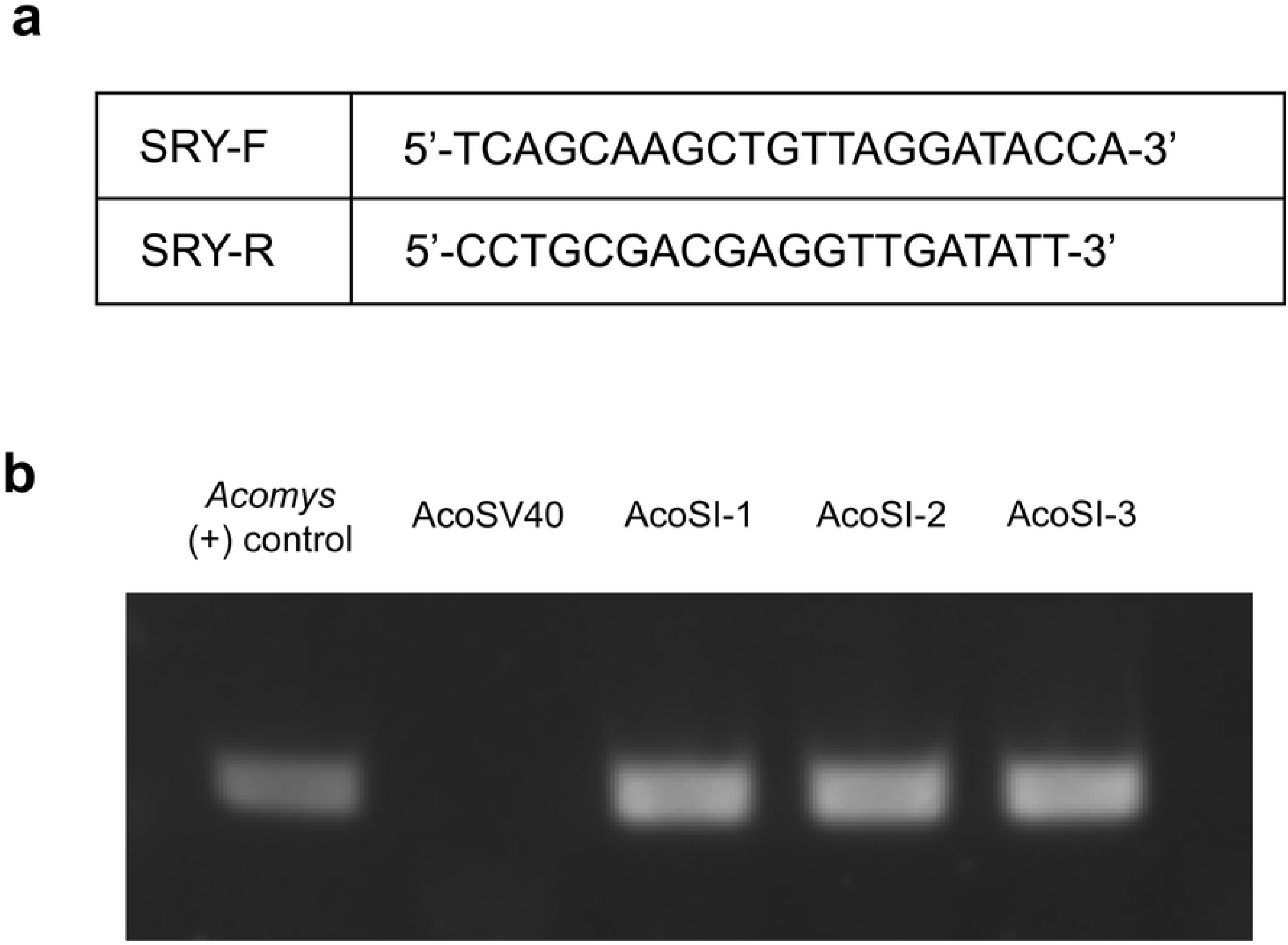
AcoSV40 cells are female while all AcoSI cells are male. (a) Forward and reverse primers specific to Acomys SrY. (b) PCR products for Y chromosome in all 4 cell lines compared to a DNA sample obtained from the blood of an *Acomys* adult male.

## Discussion

Tissue damage in humans generally results in the formation of scar tissue, which has structural and mechanical properties that differ from uninjured tissue and often impede tissue function. Regenerative organisms present an opportunity to uncover mechanisms behind scar-free healing that can improve patient outcomes following tissue damage. The spiny mouse (*Acomys*) is an exciting research organism with the most extensive regeneration capabilities of any known mammal. Unfortunately, *Acomys* research is currently limited to a handful of institutions due to the need for non-traditional animal facilities and husbandry protocols, as well as the limited access to *Acomys* vendors [1,18–20]. To increase access to *Acomys* research and reduce the use of animals in regenerative medicine research, we developed two immortalized *Acomys* fibroblast cell lines. We generated the lines through two well described methods— SV40LT transfection (“AcoSV40”) and spontaneous immortalization (“AcoSI-1”)—and assessed morphological and functional characteristics.

As mediators of matrix deposition and wound contraction, fibroblasts are likely key players in the scar-free healing of *Acomys. Acomys* wounds have low populations of αSMA positive myofibroblasts [2], even though there is a high concentration of TGF-β1 [3], a pro-fibrotic cytokine. *In vitro, Acomys* fibroblasts do not upregulate αSMA when treated with TGF-β1 [50]. These findings suggest that *Acomys* fibroblasts are protected from TGF-β1-mediated activation. Inhibition of TGF-β1 signaling *in vivo* is associated with reduced wound scarring, as demonstrated in a rabbit ear hypertrophic scarring model [53]. However, direct blocking of TGF-β through antibody-based methods have been unsuccessful due to adverse effects [54]. Uncovering mechanisms behind the response of *Acomys* fibroblasts to TGF-β1 may lead to more nuanced treatments targeting the TGF-β pathway. In addition to an altered response to TGF-β1, *Acomys* fibroblasts demonstrate an increased migration rate and decreased response to stiffness *in vitro*, which may also play roles in wound healing and fibrosis, respectively [45]. Proliferation and migration of dermal fibroblasts is a crucial step in wound healing due to the role of fibroblasts in depositing granulation tissue and beginning the proliferative phase of healing [55]. The lack of stiffness-induced myofibroblast differentiation in *Acomys* fibroblasts is a deviation from the common feed-forward mechanism between increased ECM deposition and myofibroblast activation that occurs in fibrotic disorders [56]. Understanding these intrinsic differences between *Acomys* fibroblasts and fibroblasts in non-regenerative mammals may point to new therapeutics to improve wound healing and mediate fibrosis.

AcoSV40 and AcoSI-1 immortalized *Acomys* fibroblasts present a great opportunity for researchers to uncover new mechanisms behind scar-free wound healing. The two cell lines are easier to culture than pAFs due to higher proliferation rates, consistent characteristics, and simplified media requirements (S3 Fig). Their off-the-shelf availability allows for more frequent experiments and collaboration across institutions. Although the two cell lines behave similarly, there are instances where one cell type may be preferred over the other. For example, AcoSV40 cells express puromycin N-acetyltransferase, a puromycin resistance gene commonly used in selection of transformed mammalian cells. This prevents the use of puromycin selection in any further genetic modifications of these cells, making the AcoSI-1 cells preferable if using lentiviral vectors with puromycin resistance. In addition, AcoSI-1 cells may be a more appropriate comparison when compared to NIH3T3 fibroblasts because they are both spontaneously immortalized lines and have low αSMA expression in standard culture conditions. However, if αSMA expression is desired under standard culture conditions, AcoSV40 fibroblasts would be preferred over AcoSI-1. Like αSMA, other proteins may be differentially expressed between the two cell lines and a decision between the two lines will need to be made based on the proteins of interest for specific experiments. Finally, AcoSV40 cells are female and AcoSI-1 cells are male, which may affect cell line choice depending on research questions, as there are differences between male and female cells from a variety of tissues [52].

To increase access to *Acomys* research and reduce the use of animals in regenerative medicine research, we developed two immortalized *Acomys* fibroblast cell lines and confirmed that morphological and functional characteristics were representative of pAFs. We believe the availability of these cells will contribute to the field’s understanding of mammalian regeneration.

## Acknowledgements

The authors gratefully acknowledge assistance from the Mass Spectrometry Research and Education Center with proteomic analysis and ALSTEM Inc. for immortalization services related to the AcoSV40 fibroblasts. The authors would also like to acknowledge Malcolm Maden for his assistance with isolation of primary *Acomys* and *Mus* fibroblasts, Janak Gaire for assistance with PCR, and Paulina W. Kapuscinska for her work on Supplemental Figure 2.

## Supporting Information

**Supplemental Figure 1. Confirmation of expression of SV40 and puromycin resistance gene.** (a) Primer sequences used to analyze transgene expression, provided by ALSTEM, Inc. (b) PCR products to confirm transgene expression in AcoSV40 cells. Lane 1. MEF Sample for SV40; Lanes 2 and 5, ladder; Lane 3 and 6, positive control; Lane 4, MEF Sample for puromycin. After amplifying with primers SV40-F/R and puro-F/R, respectively, the MEF cells showed 112 bp bands for SV40 and 198 bp bands for puromycin resistance gene.

**Supplemental Figure 2. AcoSV40 and AcoSI-1 fibroblasts maintain constant proliferation rates when cultured in a standard media formulation (DMEM, 10%FBS, 1%PenStrep).** Fibroblasts were seeded at a constant cell density (5,300 cells/cm^2^) and passaged every 3 days.

**Supplemental Figure 3. Primary *Acomys* fibroblasts demonstrate faster proliferation compared to primary *Mus* fibroblasts.** Growth curves for early passage primary *Acomys* (blue) and *Mus musculus* (orange) fibroblasts. Fibroblasts were seeded at a constant cell density (5,300 cells/cm^2^) and passaged at 80-90% confluency.

